# Prediction of complex phenotypes using the *Drosophila* metabolome

**DOI:** 10.1101/2020.06.11.145623

**Authors:** Palle Duun Rohde, Torsten Nygaard Kristensen, Pernille Sarup, Joaquin Muñoz, Anders Malmendal

## Abstract

Understanding the genotype – phenotype map and how variation at different levels of biological organization are associated are central topics in modern biology. Fast developments in sequencing technologies and other molecular omic tools enable researchers to obtain detailed information on variation at DNA level and on intermediate endophenotypes; such as RNA, proteins and metabolites. This can facilitate our understanding of the link between genotypes and molecular and functional organismal phenotypes. Here, we use the *Drosophila* Genetic Reference Panel and nuclear magnetic resonance (NMR) metabolomics to investigate the ability of the metabolome to predict organismal phenotypes. We performed NMR metabolomics on four replicate pools of male flies from each of 170 different isogenic lines. Our results show that metabolite profiles are variable among the investigated lines and that this variation is highly heritable. Secondly, we identify genes associated with metabolome variation. Thirdly, using the metabolome gave better prediction accuracies than genomic information for four of five quantitative traits analysed. Our comprehensive characterization of population-scale diversity of metabolomes and its genetic basis illustrates that metabolites have large potential as predictors of organismal phenotypes. This finding is of great importance e.g. in human medicine and animal and plant breeding.

## 1. Introduction

Understanding how information encoded in the DNA is transcribed to RNA, translated to proteins and other downstream endophenotypes such as metabolites, and how this information dictates the organismal functional phenotype is in the core of several biological disciplines. While the revolutionary work that led to the discovery of this central flow of genetic information within biological systems was published more than 70 years ago (Crick 1970) we are still making progress in understanding the genotype – phenotype map. This is aided by new technologies within molecular and systems biology, allowing researchers to obtain full genome sequences from individuals of any species, detailed information about expression levels of all genes, abundancies of proteins and metabolites etc. These omics tools have provided unforeseen knowledge about the genetic and environmental background of complex traits, which has revolutionized the field of genetics with strong impacts on multiple research disciplines including medicine, animal and plant breeding and evolutionary biology (Pinu et al. 2019; Hasin et al. 2017; Elmer 2016; Dekkers 2012).

One common goal of studies on genotype – phenotype associations is to understand to what extent a complex organismal phenotype such as behavioural traits, traits linked to reproduction, diseases or the ability to cope with stressful environmental conditions, can be predicted from DNA sequence information or from endophenotypes. If information on endophenotypes, such as transcript, protein or metabolite, can accurately predict the phenotype this has wide ranging applications across life sciences, and this has proven useful in several cases (Buckler et al. 2009; Hayes and Goddard 2010; Desta and Ortiz 2014; Hickey et al. 2017; Grinberg et al. 2019). However, studies have also demonstrated that predicting complex phenotypes based on genetic information can often be difficult and the predictive power in such studies is typically low (Schrodi et al. 2014; Sun et al. 2016; Märtens et al. 2016). Reasons for the difficulties to accurately predict the phenotype are many and include; (i) that quantitative trait values result from a complex interplay between a large number of genes, each with a small contribution to the phenotype, combined with environmental factors, (ii) despite progress, the underlying genetic architectures of most traits of medical interest or traits with relevance in agriculture or evolutionary biology are still not well-understood, (iii) the effect of so called candidate genes is often depending on the genetic background (epistasis) and they explain only a small proportion of the heritability, and (iv) genes and environments interact in their effect on the phenotype. These factors give rise to substantial challenges in constructing and implementing genetic risk prediction models across biological disciplines.

Challenges with using DNA sequence variation to predict variation at the organismal phenotypic level have sparked an interest in using endophenotypes as predictors of complex functional phenotypes (Scoriels et al. 2015; te Pas et al. 2017; Hayes et al. 2017; Van Der Ende et al. 2018). Endophenotypes influence and regulate the functional phenotype and in contrast to the genotype, which is fixed in an individual’s lifetime, they are governed by interactions between the genome of an individual and internal and external influences that range from the cellular level and to the wealth of external biotic and abiotic factors an individual is exposed to in its environment. Thus, endophenotypes have been proposed to constitute proximal links between variation at the genome level and the organismal phenotype and they may provide more accurate predictors of the functional phenotype compared to the genotype (te Pas et al. 2017; Zhou et al. 2020).

We hypothesize, that the predictive value of endophenotypes for organismal functional phenotypes is linked to the proximity of the endophenotype to the organismal phenotype. Accepting this premise information on the metabolome level should provide more accurate prediction than e.g. information at the gene-expression or protein levels. This is an emerging research field and we do not have many results on this yet. However, there are studies that support the hypothesis that transcriptomic or proteomic data combined with genotype information improves prediction of several traits in e.g. *Drosophila melanogaster* and maize (Wang and Marcotte 2010; Harel et al. 2019; Azodi et al. 2020; Li et al. 2019). The metabolome is closer to the functional phenotype than transcriptomics and proteomics and a recent study based on investigating 453 metabolites in 40 isogenic lines suggest that the metabolome might constitute a reliable predictor of organismal phenotypes and that the metabolome provide novel insights into the underpinnings of complex traits and its genetic basis (Zhou et al. 2020).

Here we elaborate on the findings from Zhou et al. (2020) by using nuclear magnetic resonance (NMR) metabolomics to obtain information about fly metabolomes in pooled samples of whole male flies from 170 inbred lines from the *Drosophila* Genetic Reference Panel (DGRP); a system of fully inbred sequenced lines of *D. melanogaster* (Huang et al. 2014; Mackay et al. 2012). NMR metabolomics constitute a highly reproducible technique that in contrast to mass spectrometry allows for metabolic profiling of the total complement of metabolites in a sample (Emwas 2015). With this set-up, we first investigated to what extent the metabolome varies across the investigated DGRP lines and whether this variation was heritable. Secondly, we performed a genome-wide scan to detect DNA sequence variants that were associated with variation in NMR feature intensity. Finally, we investigated to what degree metabolomic data could increase prediction accuracy of five complex behavioural and stress tolerance phenotypes compared to when predictions were based on DNA sequence variation.

## 2. Materials and Methods

### 2.1 *Drosophila* lines, husbandry and collection

We used 170 inbred lines of the DGRP (Mackay et al. 2012; Huang et al. 2014). The DGRP lines were established by 20 consecutive generations of full sibling inbreeding from isofemales collected at the farmer’s market in Raleigh, NC. Complete genome sequence of the DGRP lines have been obtained using Illumina platform, and is publicly available (Mackay et al. 2012; Huang et al. 2014).

The DGRP lines were maintained on standard *Drosophila* medium (see Kristensen et al. (2016) for recipe details) in a 23 °C climate chamber at 50% relative humidity with a 12h:12-h light:dark cycle. For this study only male flies, which were separated by sex after 24 hours, were used. Flies were collected from 170 DGRP lines in four replicates containing 20 flies per replicate.

### 2.2 *Drosophila* sample preparation

We prepared four replicates of 20 pooled male flies for NMR spectroscopy. Males were snap frozen at three days of age and kept at –80°C. Samples were mechanically homogenized with a Kinematica, Pt 1200 (Buch & Holm A/S, Herlev, Denmark) in 1 mL of ice-cold acetonitrile (50%) for 45 s. Hereafter samples were centrifuged (10,000 g) for 10 min at 4°C and the supernatant (900 μL) was transferred to new tubes, snap frozen and stored at –80°C. The supernatant was lyophilized and stored at −80°C. Immediately before NMR measurements, samples were rehydrated in 200 mL of 50 mM phosphate buffer (pH 7.4) in D2O, and 180 μL was transferred to a 3 mm NMR tube. The buffer contained 50 mg/L of the chemical shift reference 2,2-Dimethyl-2-silapentane-5-sulfonate-d6, sodium salt (DSS), and 50 mg/L of sodium azide to prevent bacterial growth.

### 2.3 NMR experiments and spectral processing

NMR measurements were performed at 25°C on a Bruker Avance III HD 800 spectrometer (Bruker Biospin, Rheinstetten, Germany), operating at a ^1^H frequency of 799.87 MHz, and equipped with a 3 mm TCI cold probe. ^1^H NMR spectra were acquired using a standard NOESYPR1D experiment with a 100 ms delay. A total of 128 transients of 32 K data points spanning a spectral width of 20 ppm were collected.

The spectra were processed using Topspin (Bruker Biospin, Rheinstetten, Germany). An exponential line-broadening of 0.3 Hz was applied to the free-induction decay prior to Fourier transformation. All spectra were referenced to the DSS signal at 0 ppm, manually phased and baseline corrected. The spectra were aligned using *i*coshift.(Savorani et al. 2010) The region around the residual water signal (4.87-4.70 ppm) was removed in order for the water signal not to interfere with the analysis. The high- and low-field ends of the spectrum, where no signals except the reference signal from DSS appear, were also removed (i.e., leaving data between 9.7 and 0.7 ppm). The spectra were normalized by probabilistic quotient area normalization (Dieterle et al. 2006).

Metabolite assignments were done based on chemical shifts only, using earlier assignments and spectral databases previously described (Schou et al. 2017; Malmendal et al. 2006).

### 2.4 Quantitative genetics of NMR intensities

Each NMR feature (14,440 in total) was treated as a quantitative trait. For each NMR feature, we fitted a linear mixed model to partition the total phenotypic variation (i.e. one NMR feature) into genetic and environmental variation. Using the R package qgg (Rohde et al. 2020) we fitted the model,

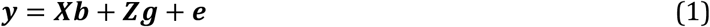

where ***y*** was a vector of NMR intensities for a particular NMR feature, ***X*** and ***Z*** are design matrices linking fixed and random effects to the phenotype, ***b*** is a vector of the fixed effects (*Wolbachia* infection status, and major polymorphic inversions; *In2Lt, In2RNS, In2RY1, In2RY2, In3LP, In3LM, In3LY, In3RP, In3RK, In3RMo, In3RC*. Information available at http://dgrp2.gnets.ncsu.edu), ***g*** is a vector of the random genetic effects defined as 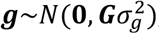, and ***e*** is a vector of residual effects defined as 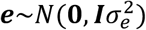. The variance structure among the random effects are modelled as independent for the residual effects (by the identity matrix ***I***), and for the genetic effects by the additive genomic relationship matrix ***G***, which was constructed using all SNPs (minor allele frequency ≥0.05) as ***G*** = ***WW***′/*m*, where *m* is the number of SNPs (i.e. 1,725,755), and ***W*** is a centered and scaled genotype matrix, where each column vector is 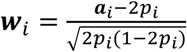, *p*_*i*_ is the allele frequency of the *i*th SNP, and ***a***_*i*_ is the *i*th column vector of the allele count matrix, ***A***, which contains the genotypes coded as 0 or 2 counting the number of the minor allele (genotypes are available at http://dgrp2.gnets.ncsu.edu).

For each NMR feature we estimated the proportion of phenotypic variation explained by SNP variation as 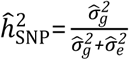, where 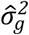 and 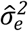 are the estimated variance components from equation 1. The significance of each 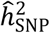 was determined as 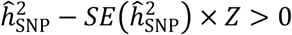, where *Z* is the quantile function of the normal distribution at probability *P* = 0.05/14,440. Thus, the resulting set of heritability estimates are those estimates that differ significantly from zero when accounting for a total 14,440 statistical tests.

### 2.5 Mapping of metabolome QTL

Metabolome quantitative trait loci (mQTLs) for mean NMR intensity were identified using linear regression (Huang et al. 2015) using the function for single marker association analysis implemented in the qgg package (Rohde et al. 2020). The estimated genetic effects (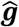, from equation 1) were used as line means, since these values represents the within DGRP line mean intensity of a single NMR feature adjusted for *Wolbachia*, chromosomal inversions, and polygenicity, which then was regressed on marker genotypes.

Significant mQTLs were identified as those SNP-NMR associations with a Bonferroni adjusted *P* < 0.05/1,725,755 = 2.9 × 10^−8^; i.e. genome-wide significant. Furthermore, we restricted the search for mQTLs to the set of NMR features with a significant heritability estimate.

### 2.6 Phenotypic predictions

Using the linear mixed model (BLUP; best linear unbiased prediction) framework we performed several phenotypic prediction models using either genomic (GBLUP) or metabolomic (MBLUP) information to investigate if the metabolome provides additional information that will increase the accuracy of prediction compared to genomic prediction. The DGRP has been characterised for a wide range of molecular, environmental stress resistance, morphological and behavioural phenotypes (Anholt and Mackay 2018; Mackay and Huang 2018). When the number of individual genotypes is low (*i.e.*, the number of DGRP lines) a large number of within-line replicates are required for accurate predictions (Edwards et al. 2016). Therefore, we restricted the set of quantitative traits to those where we had access to all individual observations, and where the average number of observations within-line was >25 (Table 1).

**Table 1.** Quantitative traits used in phenotypic predictions. Locomotor activity was assessed as distance moved per unit time (the trait was assessed with and without exposure to Ritalin (methylphenidate)). Chill come recovery is time to recover from a chill-induced coma. Startle response is a measured as the time taken to reach a certain distance following disturbance. Starvation resistance is assessed as time to death when deprived of nutrients.

| Trait | Number of<br>DGRP lines | Number of<br>observations* | $H^2$** | Lines for<br>predictions*** |
| --- | --- | --- | --- | --- |
| Locomotor activity (Rohde et al. 2019) | 172 | 30 | 0.43 | 169 |
| Locomotor activity w/ Ritalin (Rohde et al. 2019) | 172 | 30 | 0.45 | 169 |
| Chill coma recovery (Mackay et al. 2012) | 159 | 101 | 0.42 | 132 |
| Startle response (Mackay et al. 2012) | 166 | 40 | 0.50 | 137 |
| Starvation resistance (Mackay et al. 2012) | 197 | 49 | 0.52 | 166 |

\* Average number of individual observations per DGRP line.
\*\* Estimated broad-sense heritability
\*\*\* The number of DGRP lines that are in common between phenotype and the metabolomic data.

The five test traits were initially adjusted for experimental factors (see references in Table 1), *Wolbachia* infection status, and major polymorphic inversions (*In2Lt, In2RNS, In3RP, In3RK, In3RMo*). The adjusted phenotypic values were obtained as: 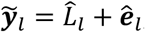, where 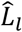 is the estimated line effect for DGRP line *l* (*i.e.*, the BLUP value), and *ê*_*l*_ is a vector containing the residuals for line *l*. Thus, ***y***_*l*_ and 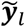 has the same dimension. The estimated line effects 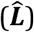 were assumed 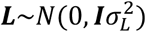. The metabolome contains aggregated information both on the individual genotypes and environmental exposures. Thus, to avoid double counting the genomic variation as represented by SNP information by adding genomic information in both the two steps in the GBLUP and MBLUP analysis, we assumed the DGRP lines to be independent, modelled by the identity matrix ***I***, instead of ***G*** in the first step.

For each quantitative trait (Table 1) we fitted two models, each for 100 randomly selected training sets (*t*, the training sets were the same for all prediction models) containing 90% of the DGRP lines:

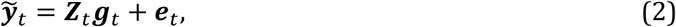

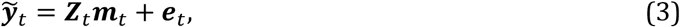

where 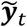 is the adjusted phenotypic values for the DGRP lines in training set *t*, ***e***_*t*_ is a vector of random residuals, ***Z***_*t*_ is a design matrix linking the genomic (***g***_*t*_) and metabolomic (***m***_*t*_) effects to the phenotypes. The random genomic effects are 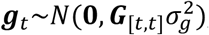, and the metabolomic effects are 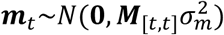, where ***G*** is the additive genomic relationship matrix as specified previously, and ***M*** is the metabolomic relationship matrix. The metabolomic relationship matrix was computed as ***M*** = ***QQ***′/*m*_NMR_, where ***Q*** is a *n* × *m*_NMR_ matrix of adjusted, centred and scaled NMR intensities (*m*_NMR_ = 14,440). Each column vector of ***Q*** contains the BLUP values from a mixed model where the phenotype was the corresponding NMR intensity, which was adjusted for *Wolbachia* infection status and major polymorphic inversions (*In2Lt, In2RNS, In3RP, In3RK, In3RMo*), using the identity matrix as covariance structure. This was done to obtain a data structure similar to the genomic data; namely one value of each DGRP line/NMR intensity.

The predicted phenotypes in validation set *v* 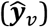 was obtained using Equation 4 and 5 for GBLUP and MBLUP, respectively.

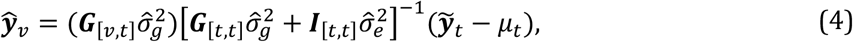

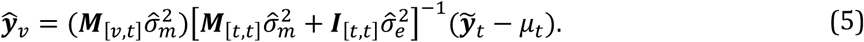

The prediction accuracy (*PA*) was quantified as the mean Pearson’s correlation between observed and predicted phenotypes across training sets, 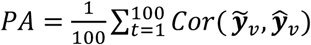. The accuracies of GBLUP and MBLUP was compared using a paired *t*-test.

### 2.7 NMR cluster-guided phenotypic predictions

From genomic prediction models we know that allowing marker effects to come from different distributions, e.g. grouping genetic variants into functional pathways, can increase the prediction accuracy markedly (Speed and Balding 2014; Edwards et al. 2016; Rohde et al. 2017, 2018; Sørensen et al. 2017; Fang et al. 2017). Therefore, we investigated if similar benefits could be achieved by partitioning the metabolome.

Using the ***Q***-matrix (i.e. the *n* × *m*_NMR_ matrix of adjusted, centred and scaled NMR intensities) we computed all pairwise Pearson’s correlation coefficients and performed hierarchical clustering on the dissimilarity on the correlation coefficients using an unweighted pair group method with arithmetic mean agglomeration (Fig. 1). Using a range of total number of clusters *K*_*cl*_ = {25, 50, 75, 100, 125, 200} we performed metabolomic feature best linear unbiased prediction (MFBLUP) which is an extension to the MBLUP model (Eq. 3) containing an additional metabolomic effect (Eq. 6, Fig. 1). For each *K*_*cl*_ level we estimated model parameters for each cluster (for *K*_*cl*_ = 25 we ran 25 models, and for *K*_*cl*_ = 50 we ran 50 models etc.) as follows:

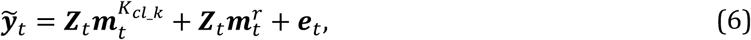

where the superscript *K*_*cl*_*k*_ indicate the total number of clusters (*K*_*cl*_) and the cluster number (*cl*). The first metabolomic effect was defined as 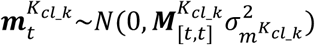, where 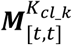 corresponds to the metabolomic relationship of the DGRP lines within the training set (*t*) for the NMR features within cluster number *cl* among the *K*_*cl*_ clusters. The second metabolomic effect 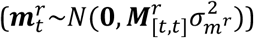 is the random effects using all NMR features except those within the *K*_*cl*_*k*_ cluster.

**Figure 1.**
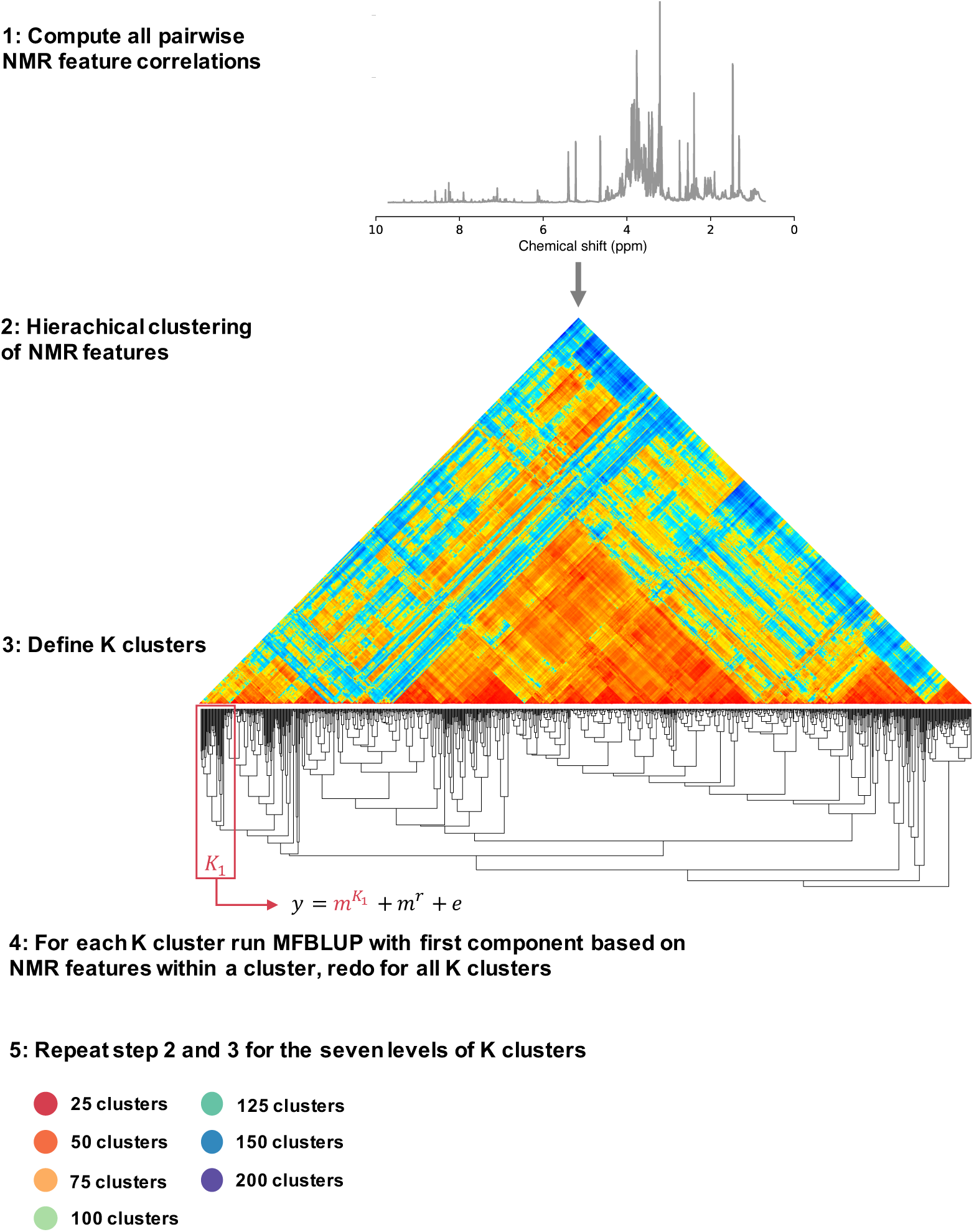
Conceptual illustration of the NMR cluster-guided phenotypic predictions. All pairwise correlations among the NMR featues were computed, which was used in a hierachical clustering of NMR features. The dendrogram was then sequentially cut into K clusters (25, 50, 75, 100, 125, 150 and 200 clusters), and each individual cluster was then used in the MFBLUP model. NMR features within one cluster was used to construct a metabolomic relationship matrix that was used as covariance matrix in the MFBLUP model. The MFBLUP model was fitted for all clusters within the seven levels of K clusters.

Similar to the MBLUP model, the predicted phenotypes in the validation set *v* 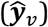 was obtained as

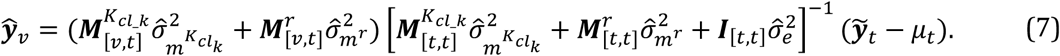

The prediction accuracy for each *K*_*cl*_*k*_ cluster was obtained as 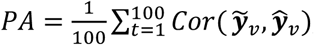, and was compared (using paired *t*-test corrected for multiple testing by a false discovery rate (FDR) of <0.05) within and across *K*_*cl*_ clusters to identify the NMR features resulting in the largest prediction accuracy. We only considered clusters to be signigicant if the FDR was below 0.05, and if the proportion of variance captured by the cluster was larger than 1%, which is computed as 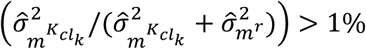.

Finaly, to investigate if we could increase the predive performance further, we took all the clusters that increased the trait-specific predictive performance (including clusters where the variance captured was below 1%), ranked them by their predictive performance, and ran a new series of prediction models, adding the NMR features sequential to the model based on the clusters predictive performance (high to low).

## 3 Results

### 3.1 The metabolome of *D. melanogaster*

Using ^1^H NMR we quantified the metabolome of males from 170 DGRP lines in four biological replicates (Supplementary Data file S1). For each of the 14,440 NMR features we estimated the proportion of variation in NMR intensity explained by common genetic variants (*h*^2^, Fig. 2A). In total, 39% of all NMR features had a significant heritability estimate (Fig. 2B), of which the average heritability was 0.26 (0.13 across all features).

**Figure 2.**
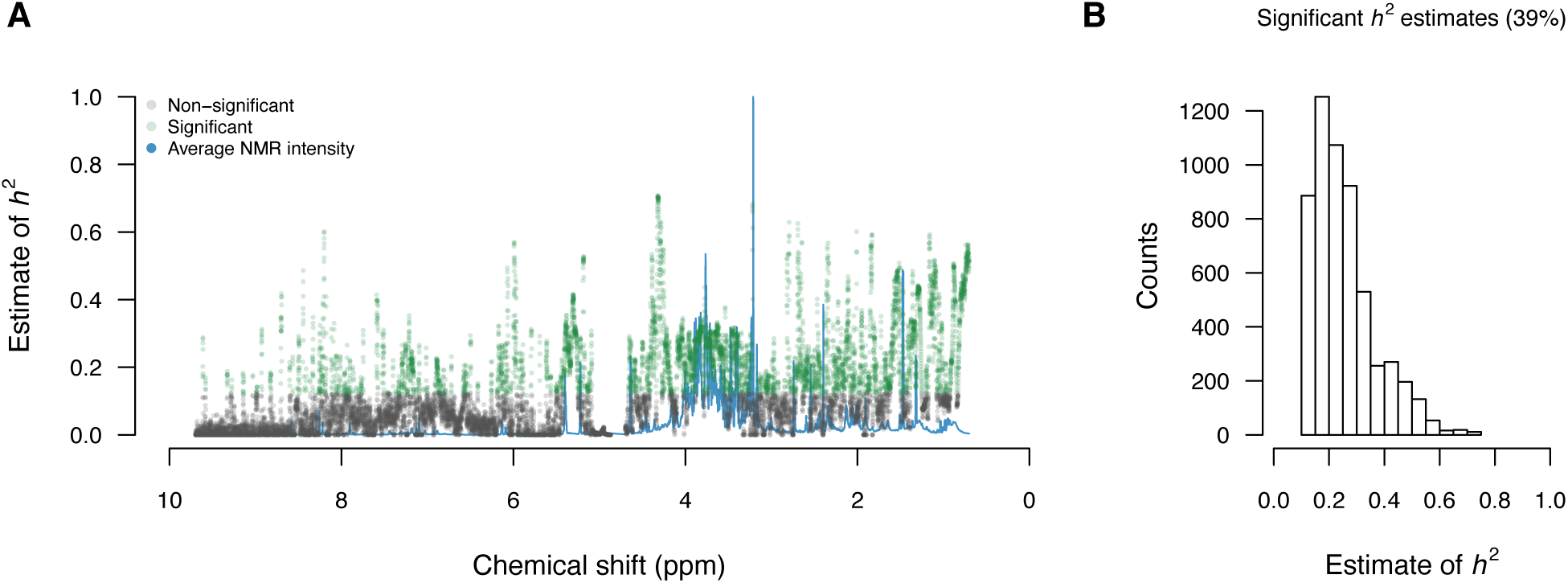
Genetic variation for the *D. melanogaster* metabolome. Panel **A** shows in solid blue line the average NMR intensity across all DGRP lines (intensity axis not shown) as function of chemical shift. For each NMR feature we estimated *h*^2^; the points in grey represent non-significant estimates of *h*^2^, and points in green are significant estimates of *h*^2^. Panel **B** is a histogram of the significant heritability estimates.

For those NMR features where we observed a significant heritability, we next sought to identify genetic variants associated with NMR feature intensity, namely mQTLs. We found a total of 5,782 genome-wide significant mQTLs (Supplementary Table 1) covering 1249 NMR features and 1468 unique genetic variants. The significant mQTLs were found within 963 genes, with the gene *Coronin* having 488 genome-wide associations (Supplementary Table S1), which is extreme compared to the average of 6.0 significant associations per gene.

### 3.2 Phenotypic predictions

To test the predictive performance of the metabolome we obtained data from five previously published data on complex phenotypes (Table 1); two behavioural traits and three stress resistance traits. We constructed relationship matrices based on genomic and on metabolomic information and performed genomic/metabolomic best linear unbiased prediction (GBLUP/MBLUP). For each trait we used 90% of the data to estimate the parameters using either the genomic or metabolomic relationship matrices and used the estimated parameters to predict the remaining 10% of the data. This was repeated on 100 random data divisions.

The mean prediction accuracy for the two behavioural traits, locomotor activity without and with treatment of Ritalin, was below 0.1 when based on genomic information (Fig. 3A-3B). Using metabolomic information the predictive performance was increased to above 0.4 (Fig. 3A-3B, Suplementary Table S2). By using the *Drosophila* metabolome, we could also increase the predictive accuracy of the two environmental stress resistance traits, chill come recovery and starvation resistance (Fig. 3C-3D, Supplementary Table S2). However, for startle response (a behavioural response to a physical disturbance) prediction using genomic information was superior over the metabolome (Fig. 3E).

**Figure 3.**
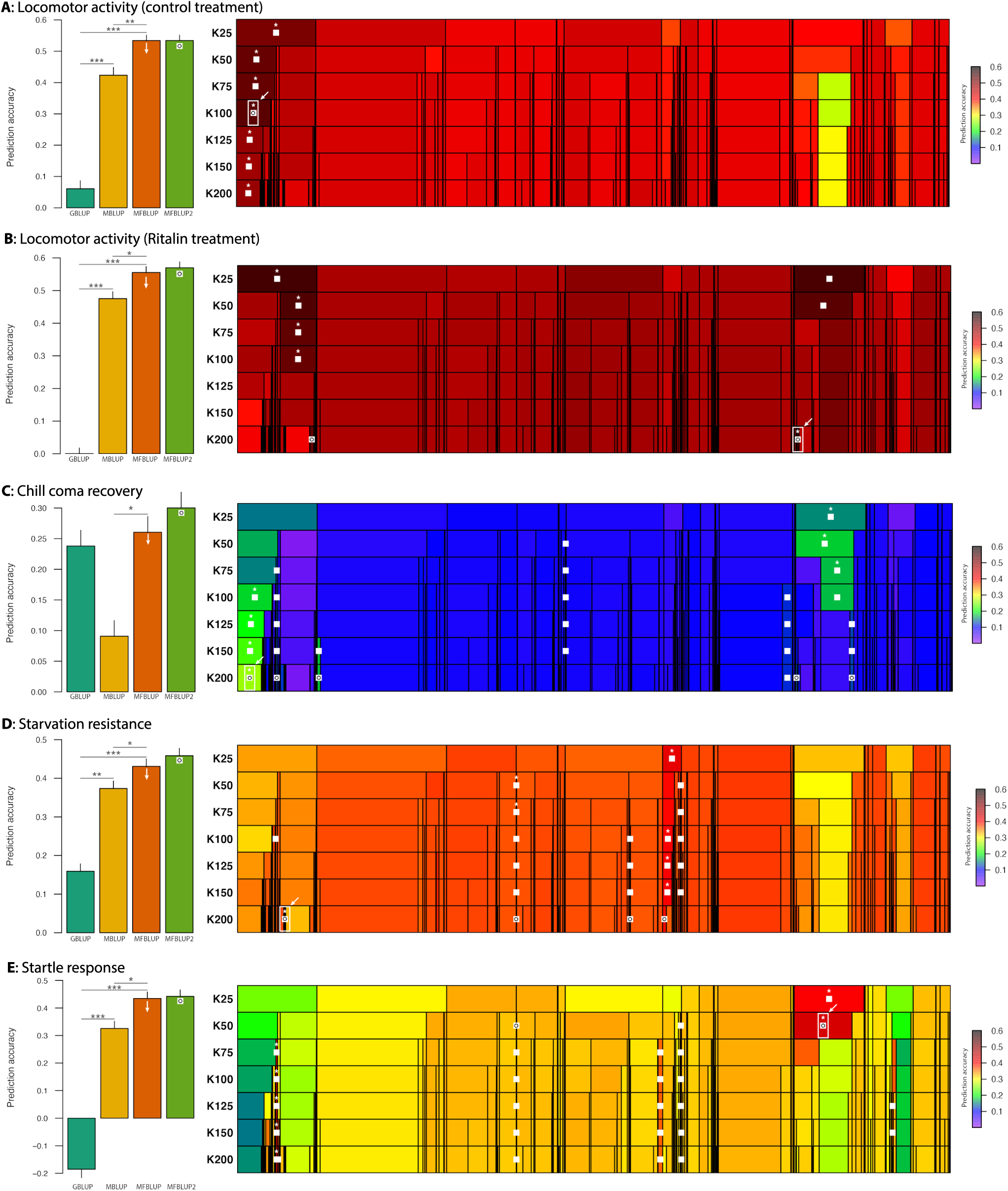
Prediction accuracies using genomic and metabolomic data. For each panel, the barplot shows the maximum mean prediction accuracies (+ standard error) for the different models. GBLUP and MBLUP are based on single component prediction models, whereas the two MFBLUP models are based on two components. The global maxium prediction accuracy obtained across all levels of clusters (*K*_*cl*_ = {25, 50, 75, 100, 125, 200}) is shown in the MFBLUP bar (indicated with white arrow). The prediction accuracy when combining the significant clusters is shown in the MFBLUP2 bar (indicated with white square and circle). Significant improved predictive performance is indicated by asterisks above the bars, see Supplementary Table S3 for all comparisons. The heatmaps on the right side of the panels show all prediction accuracies for the NMR cluster-guided prediction model within *K*_*cl*_ cluster level. The columns correspond to NMR features (fixed across the *K*_*cl*_ cluster levels) and each cell is one cluster of NMR features (link between NMR featuers and clusters can be found in Supplementary Table S4.). The predictive performance of each cluster within *K*_*cl*_ cluster level is indicated with the color scale. Within cluster level significant prediction accuracies are indicated with white squares and the cluster with the highest significant prediction accuracy is indicated with asterisk. Across all cluster levels, the highest prediction accuracy is indicated with white arrow (which then corresponds to the orange bars on the left-side panel). The set of significant clusters that combined gives the highest predictive performance is marked with white sqaures with black circle (corresponds to the light green bars in the barplot).

We computed Pearson’s correlation coefficient among all NMR features and performed hieracical clustering (Supplementary Fig. S1). We then ran the two component NMR cluster-guided prediction model (MFBLUP) where the first component was based on NMR features within one cluster, and the second component was based on the remaining NMR features (Fig. 1). We tested all clusters from the hierachical clustering using different number of total clusters; *K*_*cl*_ = {25, 50, 75, 100, 125, 200} (Fig. 1). By the extension of the MBLUP model we could increase the predictive performance of all five quantitative traits by 17%-185% (Fig. 3, Supplementary Table S2). Interestingly, the largest improvement in prediction accuracy was obtained at different cluster levels for the five traits (Fig. 3 and Supplementary Figure S2). For locomotor activity the largest improvement in prediction accuracy was obtained at cluster level *K*_*cl*_ = 100 (Fig. 3A), with cluster 1 as the only cluster that had significantly increased predictive accuracy (Supplementary Table S3 and Supplementary Fig. 3). For locomotor activity (Ritalin treatment), startle response and starvation resistance, cluster level *K*_*cl*_ = 200 contained the clusters that gave the highest predictive performance (Fig. 3B, 3D-3E and Supplementary Fig. S2). For activity (with Ritalin) cluster 121 and 74 (Supplementary Table S3) increased the predictive performance. Combining the two clusters increased the prediction accuracy insignificantly by 1% (Fig. 3B, Supplementary Figure S5). Eight clusters increased the predictive performance (that also captured >1% feature variance) for startle response (1, 2, 5, 14, 99, 112, 145 and 187, Supplementary Table S3 and Fig. S3), and by combining cluster 1, 2, 5, 112 and 145 we further increased the predition accuracy (Fig 3E, Supplementary Table S2, Supplementary Figure S7). Startle response was the only trait where metabolomic prediction performed worse than genomic prediction (Figure 3E); however, by combining the five clusters with the highest prediction accuracy the metabolomic prediction performed better than genomic prediction (Supplementary Table S2). Five clusters (21, 65, 83, 95 and 140) increased the accuracy of prediction (and captured >1% feature variance) for starvation resistance (Supplementary Fig. 3), and the joint effect of cluster 21, 65, 83 and 95 further insignificantly increased the accuracy (Fig. 3D, Supplementary Table S3, Supplementary Figure S8). Finally, for chill coma recovery the maximum prediction accuracy was obtained at cluster level *K*_*cl*_ = 50 (Fig. 3B), where cluster 5, 27 and 35 were significant explaining >1% of the NMR feature variance (Supplementary Table S3 and Supplementary Figure S3). Combining the two clusters with the highest accuracy led to insignificant increased accuracy (Fig. 3B, Supplementary Table S2 and Fig. S8).

The contributions to the predictive clusters at cluster level *K*_*cl*_ = 200 from the metabolite NMR-spectra were mapped on to the NMR spectrum (Fig. 4). Interestingly, none of the clusters contained signals from the highest concentration metabolites. Rather they contain signals from metabolites at lower abundance or very broad signals, suggesting contributions from small metabolites bound to larger molecules or larger molecules themselves. Out of the 18 clusters (Table 2), seven (1, 2, 5, 21, 32, 45, 65; Figure 4A) contained signals in a region where aromatic compunds with quartenary nitrogens, such as NAD and nicotinamide ribotide appear. These clusters showed high prediction accuracy for locomotor activity, chill coma recovery and startle response. Four clusters (32, 74, 83, 95; Figure 4A) contained signals in a region where other heterocycles, such as adenosine, appears. Furthermore, six clusters (112, 121, 129, 139, 145, 154; Figure 4B) contained signals in a region where aromatic groups from aminoacids like histidine and tyrosine appear. Most of these are important for locomotor activity with Ritalin and starvation resistance. There is also one cluster (171; Figure 4C) that contained signals in a region where signals from sugars appear, and another (187; Figure 4D) that contained signals in a region where amino acid CH2 groups appear. Out of these clusters 1 and 187 clearly contained signals that would usually be identified as baseline, while clusters 2, 5, 32, 112, 129, 139, 145 contained mostly well resolved, though often low intensity, signals. The remaining signals are somewhere in between. Only cluster 32 could be matched to a known metabolite and its signals are assigned as coming from the nicotinamide group of NADP. The other metabolites could not be found in currently available databases with NMR characteristics of metabolites (Ulrich et al. 2008; Cui et al. 2008; Wishart et al. 2018). The clusters at cluster level *K*_*cl*_ = 200 were compared with those giving the highest prediction accuracy for locomotor activity at cluster level *K*_*cl*_ = 100, and chill coma recovery at cluster level *K*_*cl*_ = 50 (Supplementary Fig. S9). For locomotor activity the larger cluster with the higher prediction activity cover a larger stretch of the baseline in the nicotinamide region indicating that it is optimal to include a larger number of higher molecular weight nicotinamide units (Fig 3A, Supplementary Fig. S9A-B). For chill coma recovery the larger cluster covers the broad peak containing the signals from aromatic amino acids in a higher molecular weight context and there is no smaller cluster that retain significant predictivity once the larger cluster is broken up (Fig. 3E, Supplementary Fig. S9C-D).

**Table 2.** Predictive NMR feature clusters.

| **Trait** | **Clusters** | **Cluster level** |
| --- | --- | --- |
| Locomotor activity | 1 | K100 |
| Locomotor activity w/ Ritalin | 74, 121 | K200 |
| Chill coma recovery | 5, 27, 35 | K50 |
| Startle response | 1, 2, 5, 14, 99, 112, 145, 187 | K200 |
| Starvation resistance | 21, 65, 83, 95, 140 | K200 |

*NMR feature IDs for each cluster can be found in Supplementary Table S4.*

**Figure 4.**
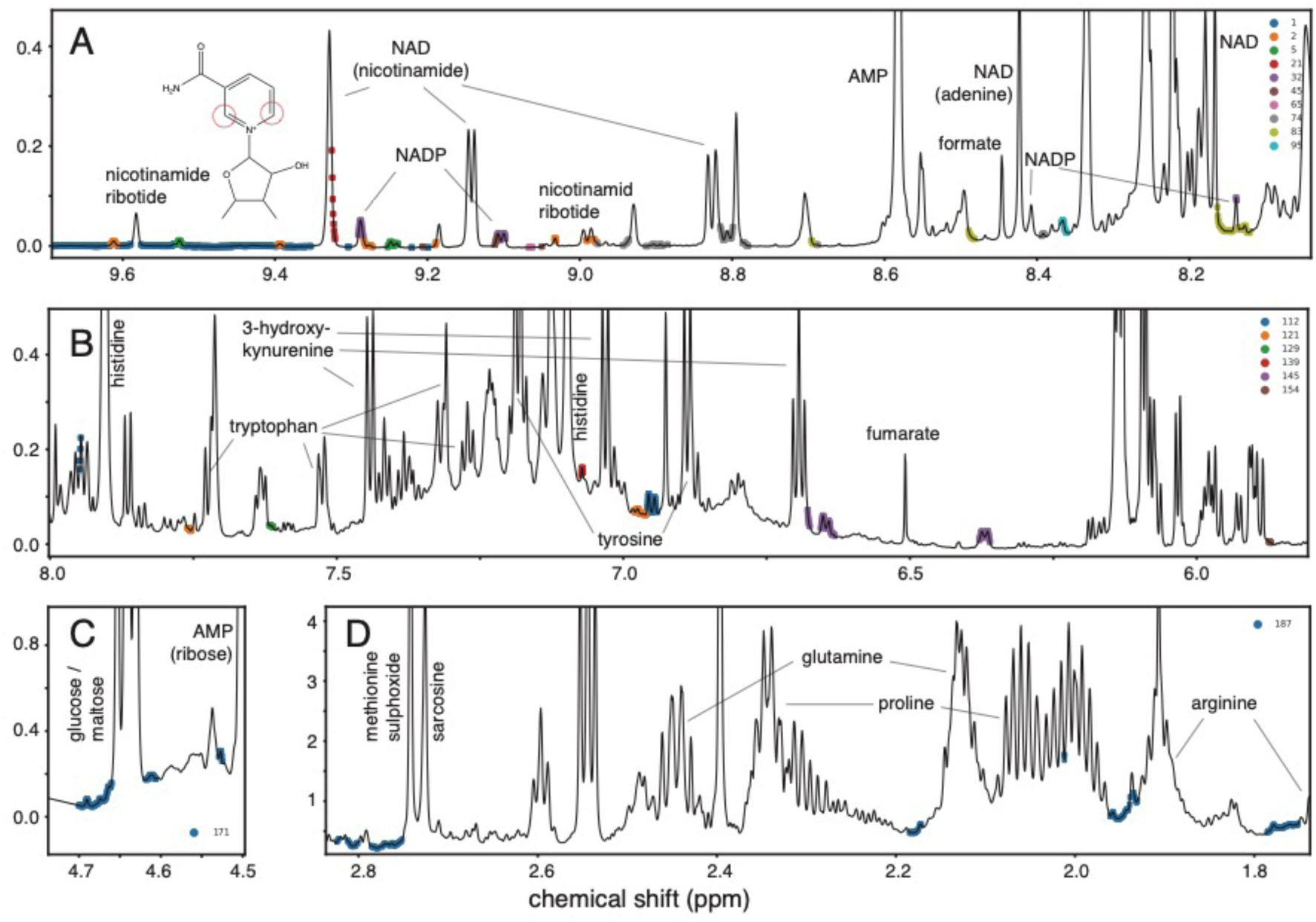
Contributions to predictive clusters from the metabolite NMR-spectra at the *K*_*cl*_ = 200 level. Clusters are indicated with colored dots on the average of all NMR spectra (black line). Panel **A** shows clusters: 1, 2, 5, 21, 32, 45, 65, 74, 83 and 95; panel **B** clusters: 112, 121, 129, 139, 145 and 154; panel **C** cluster: 171; and panel **D** cluster: 187. Selected major metabolites in these regions are identified. The location of the nicotinamide ribotide signals resonating at the highest ppm values are also indicated in panel **A**.

## 4. Discussion

Prediction of phenotypic trait values using genetic markers has been a central element in plant and livestock breeding for decades (Meuwissen et al. 2001; Van Arendonk et al. 1994; Goddard et al. 2009), and more recently this strategy has emerged within human genetics attempting to accurately predict e.g. disease risk from DNA information (Wray et al. 2008, 2019; Hall et al. 2004; Schrodi et al. 2014). However, the predictive value from genotyped genetic variants is often low (Patron et al. 2019; Schrodi et al. 2014), and this is problematic when aiming to predict complex phenotypes such as many diseases, behaviors or production traits. Therefore, there is a potential to further optimize the applicability of these methods. Here, we obtained full metabolome profiles of males from 170 DGRP lines in four biological replicates to investigate if an endophenotype, in our case the metabolome, has improved predictive power compared with a situation where only genome information is available.

We found that the *Drosophila* metabolome was highly variable with more than 39% of the NMR features having a significant heritability estimate (Fig. 2), displaying same level of genetically determined variability as the *Drosophila* transcriptome (Huang et al. 2015). It has previously been shown that metabolome variation appears to have a genetic signature (Zhou et al. 2020), but also that the metabolome is highly variable among sexes (Zhou et al. 2020; Li et al. 2018) and change with age (Hoffman et al. 2014; Yoshida et al. 2010; Lawton et al. 2008). Our findings confirm that variation in metabolite abundance is genetically controlled. The metabolome can therefore be influenced by evolutionary forces like any other phenotypic trait and this variation can be utilized e.g. in livestock and plant breeding where specific metabolites may be of interests (Gamboa-Becerra et al. 2019; Browne and Brindle 2007; Goldansaz et al. 2017).

Metabolite-wide association studies, which is the mapping of metabolite QTLs, seeks the same as genome-wide and transcriptome-wide association studies, namely to identify genetic variants associated with variation in a functional character or an endophenotype (Holmes et al. 2008; Bictash et al. 2010). Here, we mapped the individual data points in the metabolome spectra with SNP genotypes and identified abundant metabolite feature – SNP associations (Supplementary Table S1). We identified more than 900 genes associated with metabolome variation, where the average number of associations per gene was six. Such a low number of features may make it difficult to identify the underlaying metabolites. However, four genes had more than 100 feature – SNP associations. The *Drosophila* gene *Coronin* was the gene containing most significant associations with metabolome variation (488 metabolomic features). *Coronin* is involved in muscle morphogenesis (Schnorrer et al. 2010). The human orthologue of *Coronin* is *coronin 1C* and has previously been associated with lipoprotein and cholesterol levels (Siewert and Voight 2018; Wakil et al. 2016). Accordingly, the NMR features associated with *Coronin* include signals corresponging to the choline methyl groups of sn-glycerophosphocholine and methylene and methyl groups from larger molecules such as peptides or fatty acids (Supplementary Fig. S10A). The second most associated gene was *sidestep* (133 NMR features), which controls the migration of motor axons in the developing fly (Siebert et al. 2009). Across model organisms and humans *sidestep* has no apparent orthologous genes (Hu et al. 2011). Another gene with many associated NMR features (123) was *CG43373*, which has the human ortholog adenylate cyclase 5 (*ADCY5*), that in multiple studies have been associated with type II diabetes mellitus (Qi et al. 2017; Mahajan et al. 2014; Bonàs-Guarch et al. 2018), body mass index (Locke et al. 2015), blood glucose (Manning et al. 2012) and cholesterol levels (Liu et al. 2017; Hoffmann et al. 2018). Both of these genes associate with the same unidentified signal in a region with signals from hydrogen in the vicinity of hydroxy or carboxy groups (Supplementary Fig. S10B-C). These two genes have no SNP’s in common, so they are only highly correlated. The last gene with more than 100 associated NMR features is *CCHa2r* that encodes a neuropeptide (Hansen et al. 2011). In humans, *CCHa2r* is the bombesin receptor subtype 3, which regulates metabolic rate (Xiao et al. 2017) and glucose metabolism (Feng et al. 2011). This gene associates with signals from tyrosine only (Supplementary Fig. S10D). Our results clearly indicate that the human orthologues of the top associated genes are involved in metabolic processes supporting that the metabolomic feature-SNP associations are biological relevant and not statistical artefacts. The genes shortly presented here and other genes strongly associated with metabolome variation (Supplementary Table S1) are candidates for further studies using e.g. knock in and knock out technologies to verify their importance at the functional phenotypic level.

Accurate phenotypic prediction of any trait requires large sample sizes to reliably estimate model parameters, and some measure that can describe the covariance structure among individuals, such as genetic variants or other molecular variation. Although the DGRP system appears to lack power for mapping and prediction studies because of the limited number of inbred lines, it gains statistical power because it is possible to obtain repeated measures on a large number of individuals with the same genetic background resulting in very accurate within-DGRP-line measures (Mackay and Huang 2018).

The main aim of the current study was to investigate the predictive power of the *Drosophila* metabolome compared with prediction models using genomic data. We showed that for four out of five quantitative traits using the *Drosophila* metabolome for phenotypic predictions significantly improved the accuracy of prediction compared to using the *Drosophila* genome (Fig. 3). The extent to which prediction was increased varied across traits, but the trend is clear across all five traits; partitioning the metabolome by highly correlated NMR features increased the predictive performance (Fig. 3). Despite lack of significance for startle response (Fig. 3G), the increase in predictive ability was bordeline significant (*P* value = 0.09).

Recently, Zhou et al. (2020) measured metabolite variation in 453 metabolites using 40 DGRP lines, and concluded that if the sample size was larger the accuracy of metabolome predictions could be improved. This is exactly what we have shown in this study; that the metabolome can increase the accuracy of phenotypic prediction. The increased power relative to Zhou et al. (2020) may originate not only from the larger numbers of DGRP lines, but also from the higher reproducibility that NMR metabolomics provide. Moreover, we have demonstrated that our novel approach, NMR cluster-guided phenotypic prediction, identifies metabolite modules of NMR features which are trait specific (Table 2 and Fig. 3), which significantly increased the prediction accuracy compared to when using the entire metabolome (Fig. 3).

Our results clearly demonstrate the added value of performing predictions of functional phenotypes using NMR metabolomics compared with SNP genotypes. These findings, together with others, truly open the doors for applying metabolomics in different disciplines, for example in the human health sector or in animal breeding. Metabolites can be easily quantified in biofluids from livestock, human blood donors or patients that have blood samples taken on a regular basis. This entails a clear advantage over other methods in terms of translatability (Fontanesi 2016). Recently, studies have shown that the metabolomic signatures of blood from cattle (Novais et al. 2019) and pigs (Carmelo et al. 2020) can be used to accurately separate animals by their feed efficiency which is one of the most economically important traits in livestock production. Studies of the human metabolome and its relevance to human health have also clearly increased the last five years (Rangel-Huerta et al. 2019; Zhang et al. 2020), and it is very likely to be one of the cornerstones in an implementation of personalised medicine. With the emergence of large human cohort projects, like the UK Biobank (Bycroft et al. 2018), FinnGenn (Mars et al. 2020) and BioBank Japan (Nagai et al. 2017) etc., the data samples are reaching a sample level that could provide ground-breaking research if full metabolomic profiles were obtained.

The fact that many of the NMR signals with high predictivity come from unidentified and/or larger metabolites of lower concentration shows that despite the inherently low sensitivity of NMR, the high reproducibility of the method allows for a high accumulated sensitivity to be obtained for the combined data set. An interesting example here is the cluster with the highest prediction accuracy for chill coma recovery, that covers the entire aromatic amino acid region. It also shows the need for a deeper investigation of the *Drosophila* metabolome by NMR. From a metabolic signalling perspective it also makes sense that it is the less abundant metabolites with aromatic groups that are important in these processes.

In conclusion we have convincingly shown that metabolomic approaches have large potential for predicting functional phenotypes. Obviously, the generality and repeatability of these findings should be verified in different genetic backgrounds, in non-model species and in samples that are easy to generate from livestock, humans and plants. However, our main findings namely that metabolite profiles are highly heritable, that specific genes are associated with metabolome variation and that the metabolome predicts phenotypes more accurately than genomic data are robust.

## 5. Data availability

The DGRP genotypes, chromosomal inversions, *Wolbachia* infection status, and the phenotypic values for startle response, starvation resistance and chill coma recovery can be obtained from http://dgrp2.gnets.ncsu.edu/. The adjusted metabolome data can be found in the supplementary material and the locomotor activity measurements in the origical publication (Rohde et al. 2019).

## 6. Acknowledgements

The DGRP lines were obtained from the Bloomington *Drosophila* Stock Center (NIH P40OD018537, http://flystocks.bio.indiana.edu). We thank Helle Blendstrup, Susan Marie Hansen and Michael Ørsted from Aalborg University for assistance in fly maintenance and sample collection. All of the computing for this project was performed on the GenomeDK cluster. We would like to thank GenomeDK and Aarhus University for providing computational resources and support that enabled us to perform the analyses presented in the paper. The authors thank Anders Pedersen at the Swedish NMR Center at the University of Gothenburg for help with sample preparation and experimental setup and for access to the 800 MHz spectrometer. The study was supported by the Danish Natural Science Research Council through a grant to TNK (DFF-8021-00014B), and by a grant from the Lundbeck Foundation to PDR (R287-2018-735).

## Notes

### Competing Interest Statement

The authors have declared no competing interest.

### Summary of Updates

Updated Figure 3.

