## Supplementary File S1 for "Prediction of complex phenotypes using the *Drosophila* metabolome"

### Supplementary material for
#### Prediction of complex phenotypes using the *Drosophila* metabolome

Palle Duun Rohde[1,*], Torsten Nygaard Kristensen[1], Pernille Sarup[2,3], Joaquin Muñoz[1] & Anders Malmendal[4,*]

[1] *Department of Chemistry and Bioscience, Aalborg University, Aalborg, Denmark.*

[2] *Department of Molecular Biology and Genetics, Aarhus University, Tjele, Denmark*

[3] *Nordic Seed A/S, 8300 Odder, Denmark*

[4] *Department of Science and Environment, Roskilde University, Roskilde, Denmark*

---

The supplementary material contains the following:

**Supplementary Data Files**

| | |
|---|---|
| Data file S1 | *Drosophila* NMR metabolome data. [Excel file] |

**Supplementary Tables**

| | |
|---|---|
| Table S1 | Significant mQTLs. [Excel file] |
| Table S2 | Summary of prediction accuracies. |
| Table S3 | Prediction accuracies for the five quantitative traits. [Excel file] |
| Table S4 | NMR feature IDs within each cluster [Excel file] |

**Supplementary Figures**

| | |
|---|---|
| Figure S1 | NMR correlation heatmap. |
| Figure S2 | NMR-cluster guided prediction accuracies. |
| Figure S3 | Maximum prediction accuracy for NMR-cluster guided prediction. |
| Figure S4 | Aggregated cluster-guided prediction for activity, control. |
| Figure S5 | Aggregated cluster-guided prediction for activity, Ritalin. |
| Figure S6 | Aggregated cluster-guided prediction for startle response. |
| Figure S7 | Aggregated cluster-guided prediction for starvation resistance. |
| Figure S8 | Aggregated cluster-guided prediction for chill coma recovery. |
| Figure S9 | Clusters at cluster level $K_{cl} = 200$ compared with larger clusters. |
| Figure S10 | Feature maps of genes with more than 100 associated features. |

**Table S2** Summary of predictive accuracies (PA, with standard errors, SE) for prediction models based on genomic information (GBLUP) and metabolomic information (MBLUP), including model comparisons based on *t*-test on increased predictive performance.

| **Trait** | **GBLUP** | **MBLUP** | | | **Model comparison** | | | | |
|---|---|---|---|---|---|---|---|---|---|
| | **[1]** | **All NMR features [2]** | **Single cluster [3]** | **Combined clusters [4]** | **1 vs 2** | **1 vs 3** | **1 vs 4** | **2 vs 3** | **3 vs 4** |
| Activity | 0.061 (0.026) | 0.42 (0.025) | 0.53 (0.017) | 0.53 (0.017)† | *** | *** | *** | ** | - |
| Activity w/ Ritalin | 0.0004 (0.017) | 0.48 (0.022) | 0.56 (0.018) | 0.57 (0.018) | *** | *** | *** | * | ns |
| Chill coma recovery | -0.19 (0.031) | 0.33 (0.026) | 0.43 (0.023) | 0.44 (0.024) | *** | *** | *** | * | ns |
| Startle response | 0.24 (0.026) | 0.091 (0.028) | 0.26 (0.027) | 0.30 (0.025) | ns | ns | ns | * | ns |
| Starvation resistance | 0.16 (0.019) | 0.37 (0.016) | 0.43 (0.015) | 0.46 (0.015) | ** | *** | *** | * | ns |

† *set only contains one cluster, thus it is equal to the single cluster with highest predictive performance (and no P value is computed).*

*** *P value < $2.2\times10^{-16}$ for significant increase in prediction accuracy after correcting for multiple testing*

** *P value < 0.001 for significant increase in prediction accuracy after correcting for multiple testing*

* *P value < 0.05 for significant increase in prediction accuracy after correcting for multiple testing*

*ns P value > 0.05 for significant increase in prediction accuracy after correcting for multiple testing*

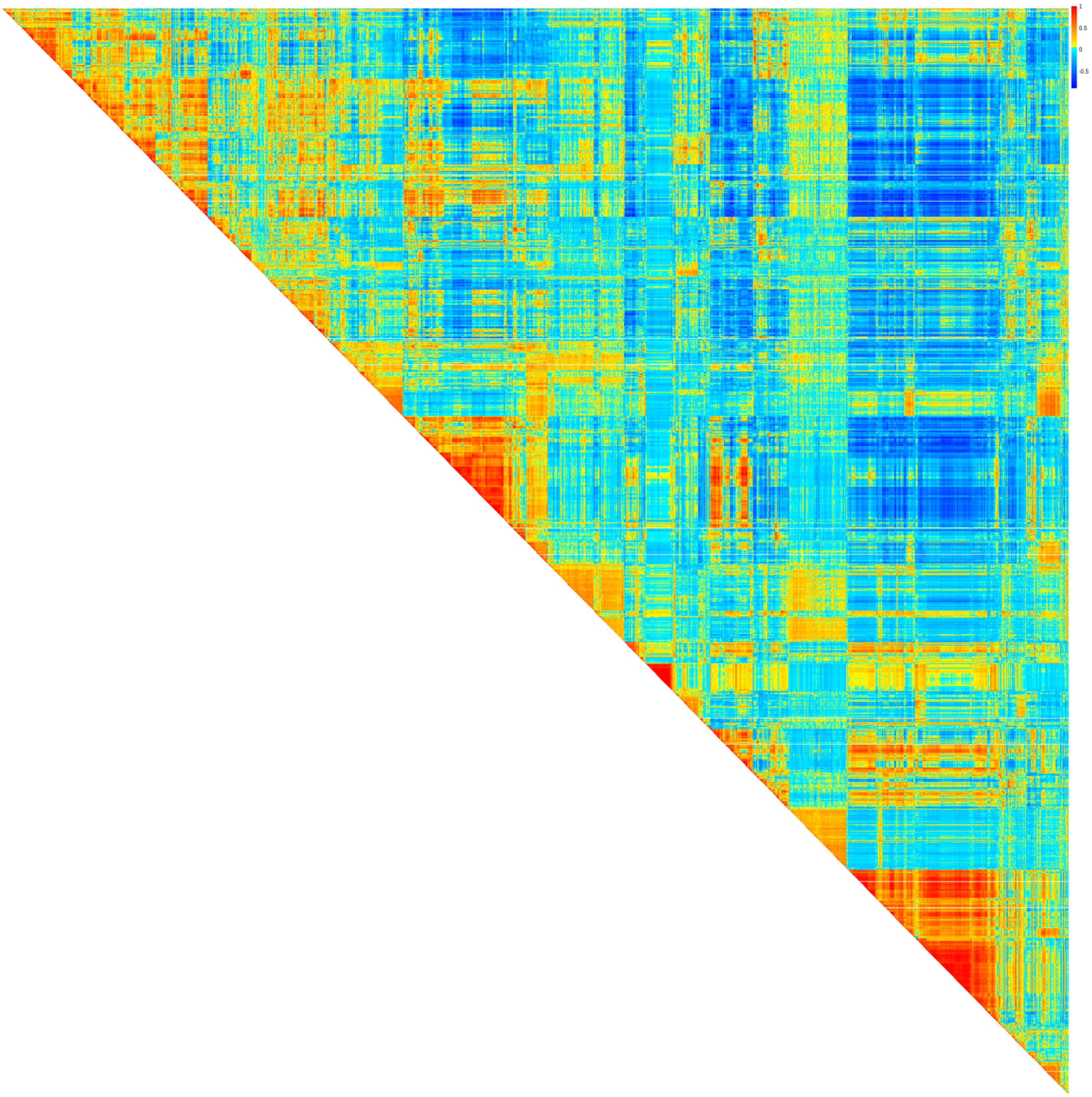


**Figure S1 Heatmap of correlations among NMR features.** Rows and columns are ordered based on hierarchical clustering.

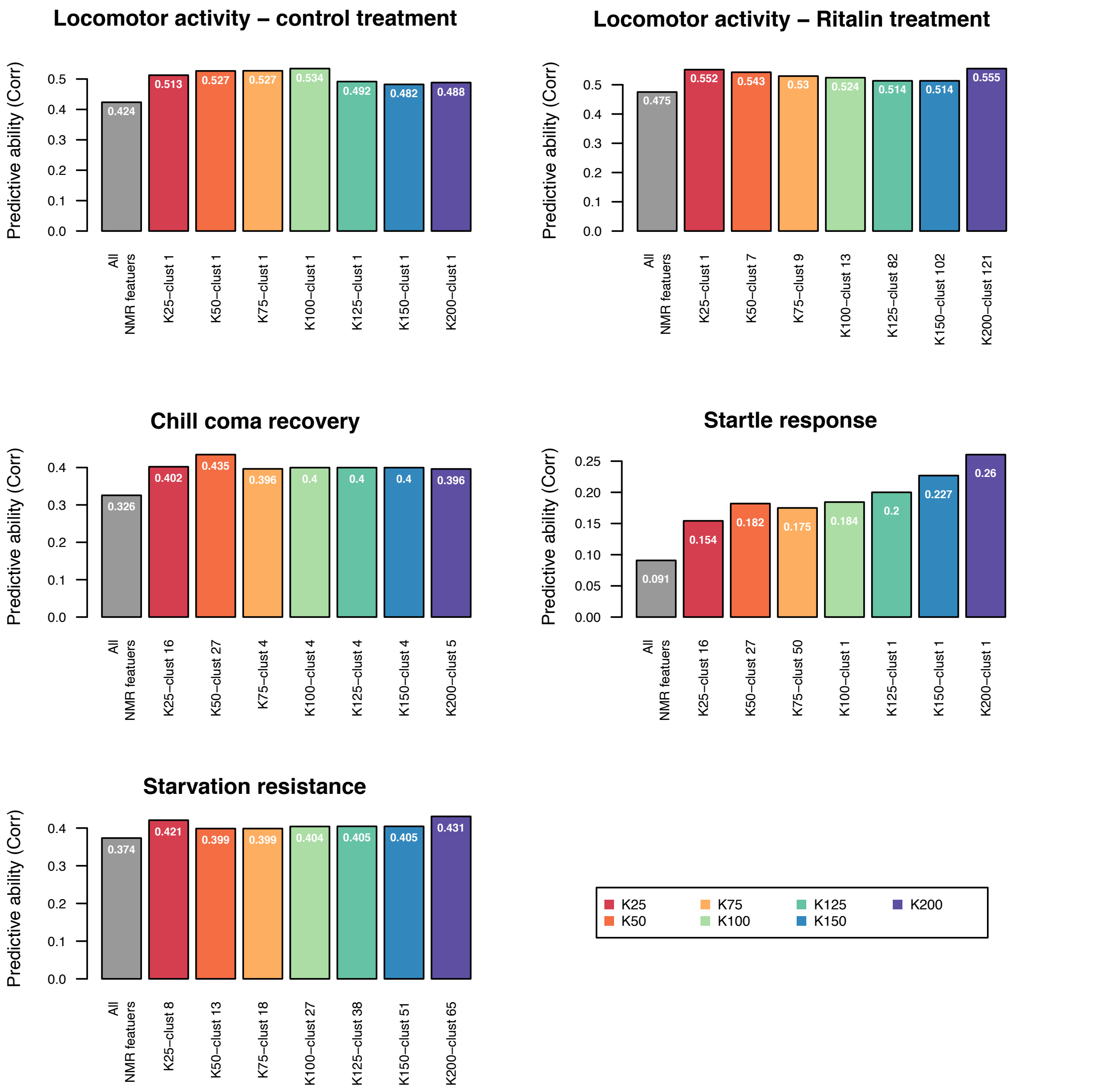


**Figure S2 Maximum prediction accuracy for NMR cluster-guided prediction models.** Each par shows the maximum prediction accuracy obtained at each cluster level $K_{cl} = \{25, 50, 75, 100, 125, 200\}$.

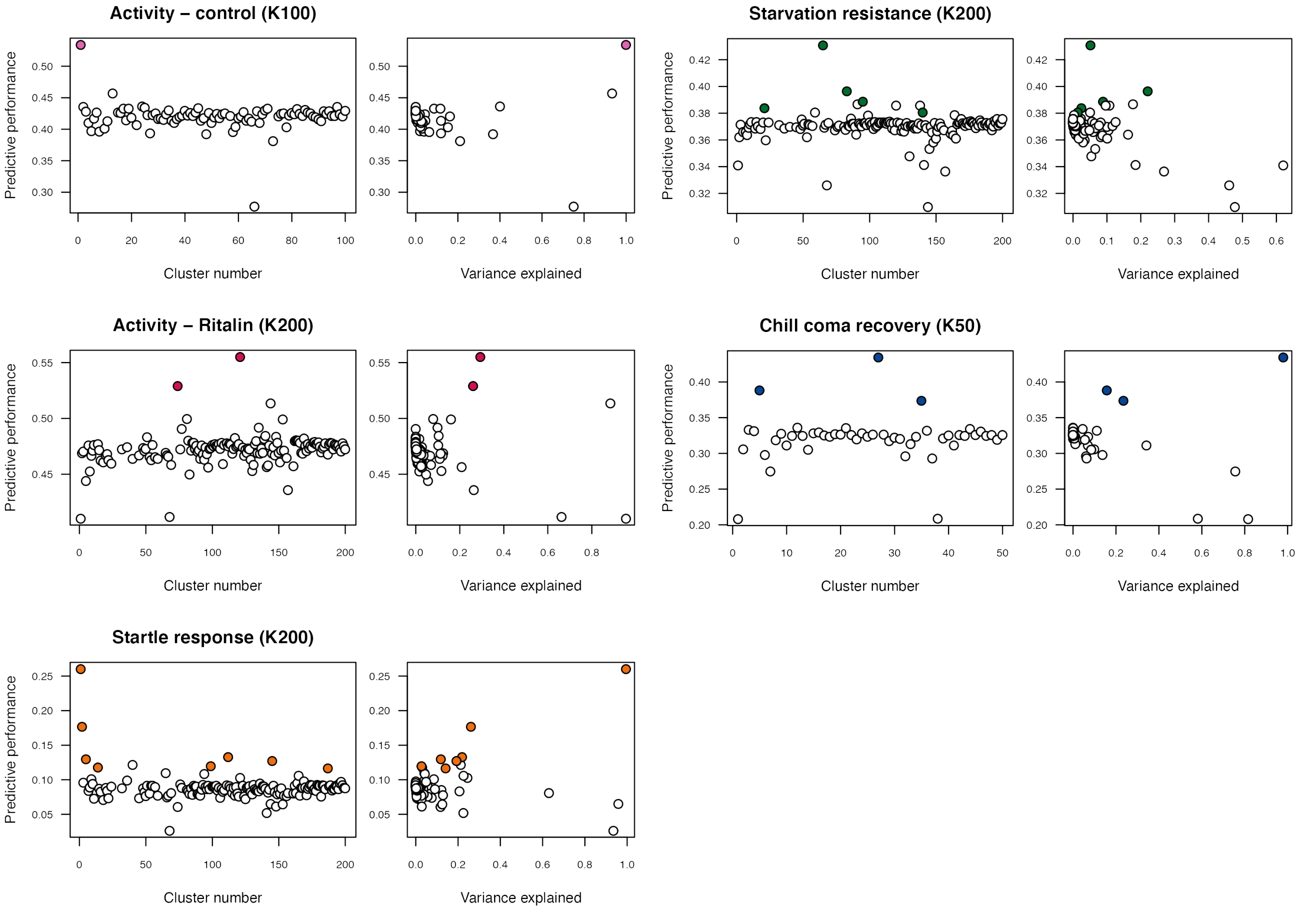


**Figure S3 Maximum prediction accuracies of the NMR cluster-guided predictions.** The largest improvement in prediction accuracy was obtained at different cluster sizes. For each trait the mean prediction accuracy (the correlation between observed and predicted phenotypes) and mean variance explained by the NMR cluster is shown for all clusters at the cluster level that gave rise to the highest prediction accuracy.

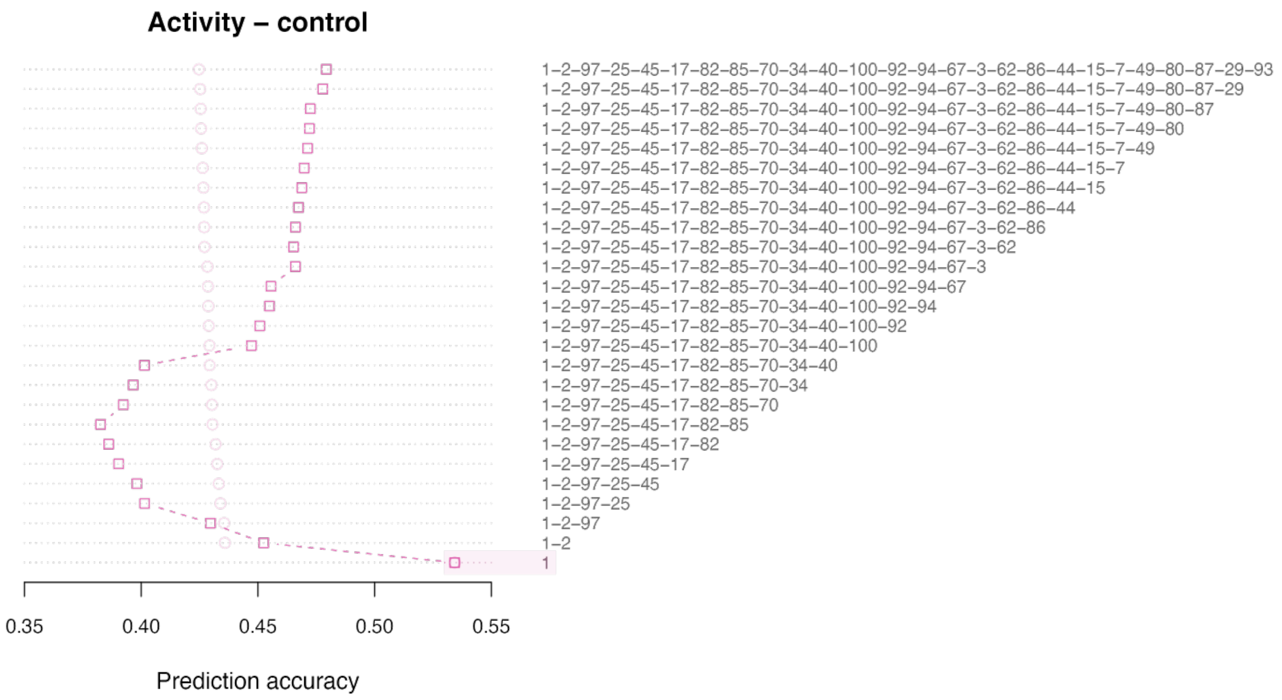


**Figure S4 Aggregated cluster-guided prediction for locomotor activity (control treatment)**.

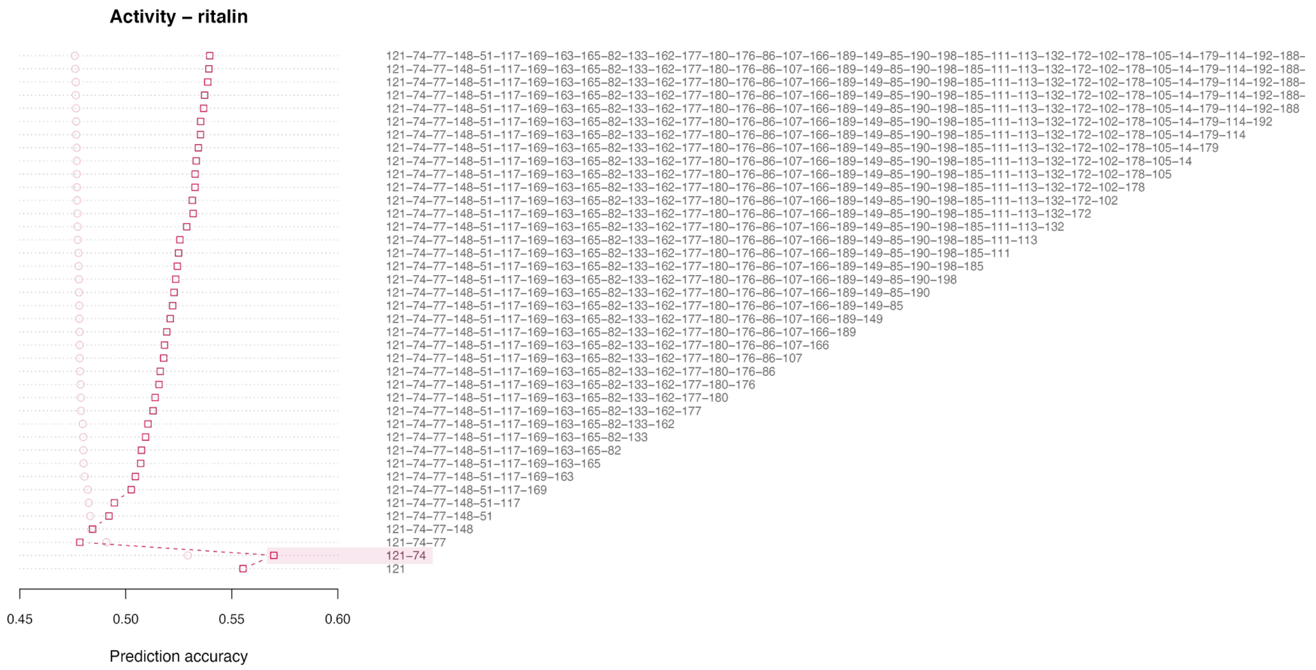


**Figure S5 Aggregated cluster-guided prediction for locomotor activity (Ritalin treatment)**.

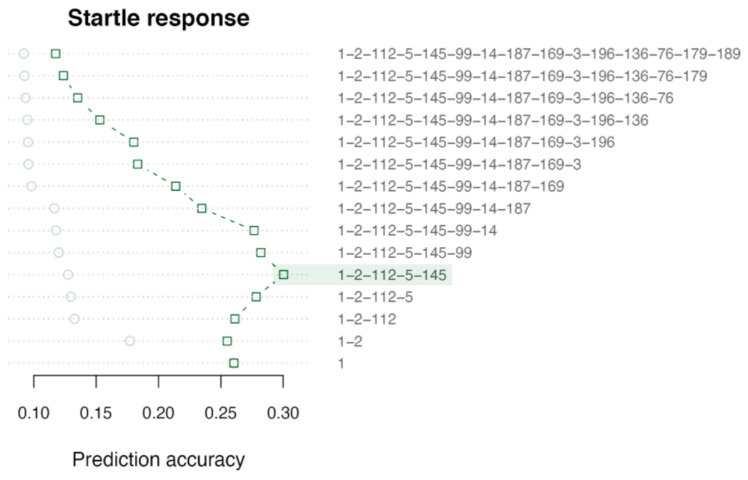


**Figure S6 Aggregated cluster-guided prediction for startle response.**

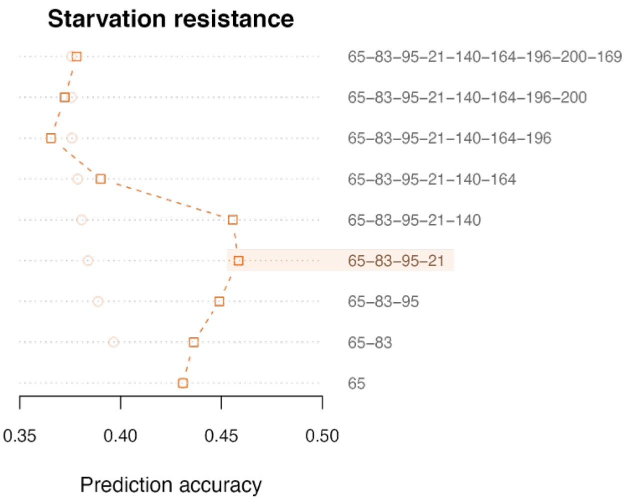


**Figure S7 Aggregated cluster-guided prediction for starvation resistance.**

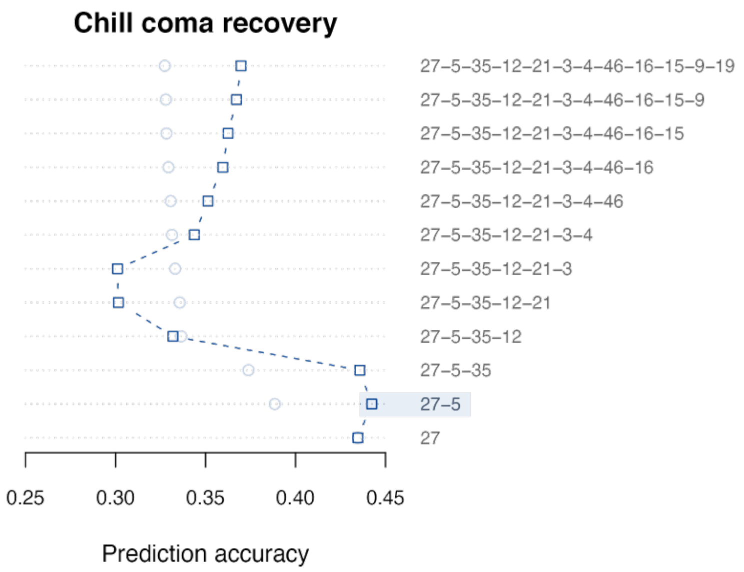


**Figure S8 Aggregated cluster-guided prediction for chill coma recovery.**

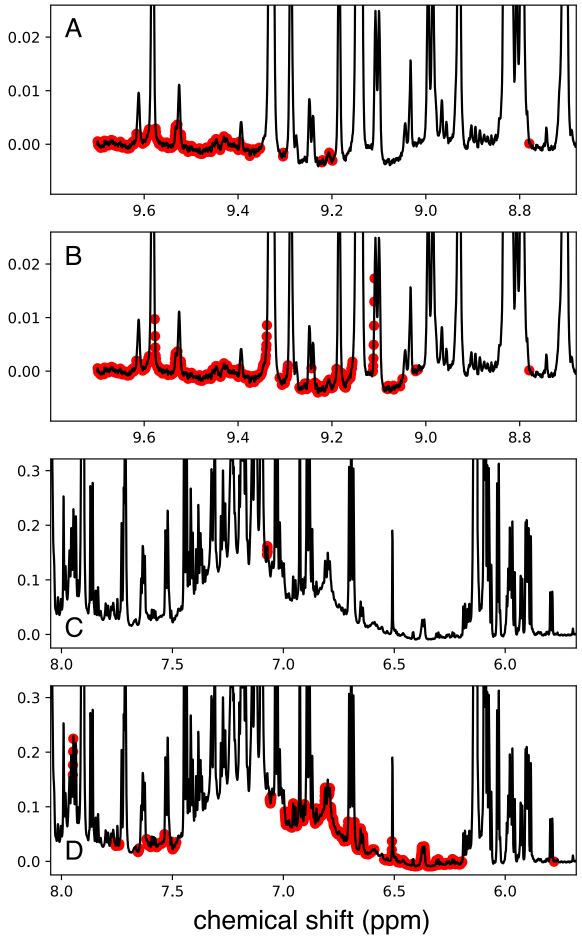


**Figure S9.** Clusters at cluster level $K_{cl} = 200$ compared with larger clusters giving the higher prediction accuracy. For locomotor activity clusters at level $K_{cl} = 200$ (A) and $K_{cl} = 100$ (B) are shown, and for chill coma recovery at clusters levels $K_{cl} = 200$ (C) and $K_{cl} = 50$ (D) are shown. Only clusters in the region covered by the larger cluster are show.

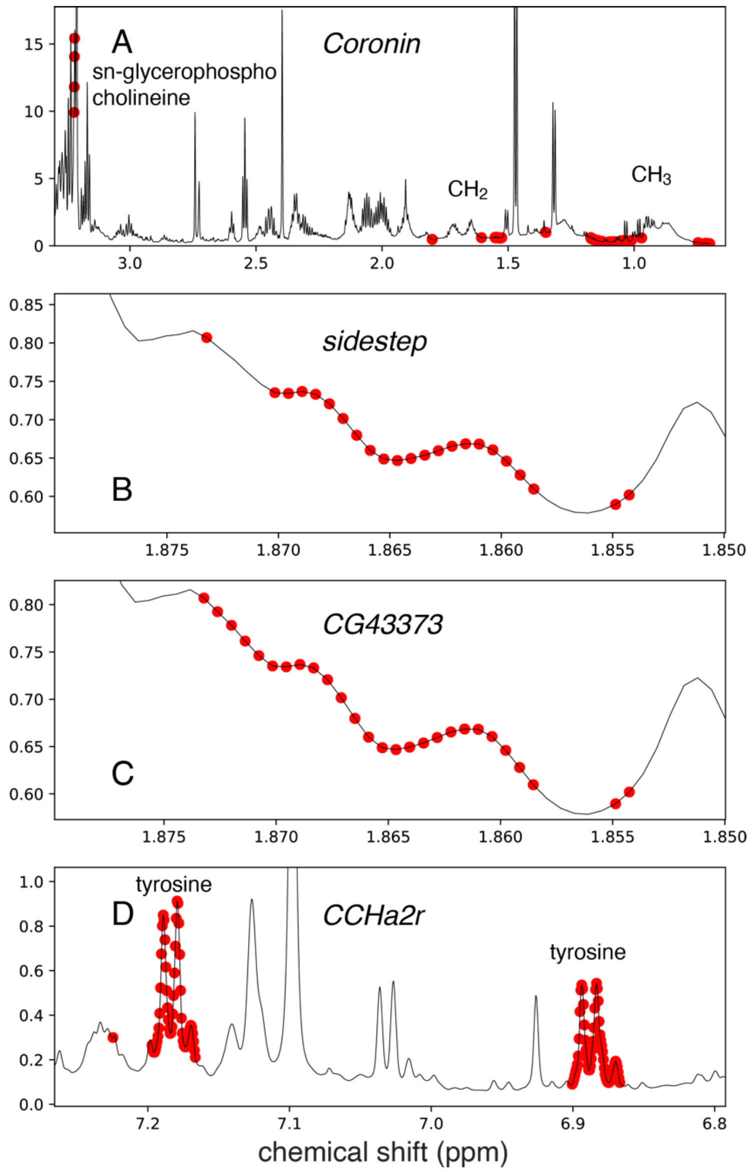


**Figure S10 Feature maps of genes with more than 100 associated features.** For A and D all features are shown, while for B and C there are also some features in other parts of the spectrum.
